# MicroED with the Falcon III direct electron detector

**DOI:** 10.1101/615484

**Authors:** Johan Hattne, Michael W. Martynowycz, Tamir Gonen

## Abstract

Microcrystal electron diffraction (MicroED) combines crystallography and electron cryomicroscopy (cryo-EM) into a method that can be used for high-resolution structure determination. In MicroED nanosized crystals, often intractable by other techniques, are probed by high-energy electrons in a transmission electron microscope and the diffracted signal is recorded on an electron detector. Since only a small number of different detectors have been used for MicroED measurements in the past, their impact on data quality has not been investigated. Here we evaluate two different cameras using crystals of the well-characterized serine protease proteinase K. Compared to previously used equipment, the Falcon III direct electron detector and the CMOS-based CetaD camera can collect complete datasets both faster and using lower total exposure. As an effect of the lower dose, radiation damage is reduced, which is confirmed in both real and reciprocal space. The increased speed and lower exposure requirements have implications on model quality and the prospects for further automation of MicroED.

## Introduction

Microcrystal electron diffraction (MicroED) is a diffraction method exploiting the strong interaction of electrons with matter to determine high-resolution structures from crystallized samples (Shi *et al.*, 2013). Owing to the favorable ratio of elastic to inelastic interactions (Henderson, 1995), MicroED can be used to collect useful data from crystals that are much smaller than required for *e.g*. X-ray crystallography. This is a significant advantage, since obtaining crystals that are both large and sufficiently well-ordered to yield high-resolution diffraction data, often constitutes a significant bottleneck in crystallography. Because crystals are imaged and screened using the same optical elements that are ultimately used to collect diffraction data, the large magnification of a transmission electron microscope (TEM) can be leveraged to select crystals with side lengths as small as 50 nm (Rodriguez *et al.*, 2015). In contrast to single-particle cryo-EM, the near-perfect alignment of the molecules in a crystal implies that the measured signal is strong enough to yield high-resolution structural information from even small peptides or chemical compounds (Jones *et al.*, 2018). MicroED thus provides a means to determine structures that are not attainable by other means.

In MicroED, crystals are continuously rotated in the electron beam of a TEM and a fast camera is used to record a shutterless movie of the resulting diffraction patterns (Nannenga *et al.*, 2014). A camera for electron diffraction data collection must have a high dynamic range such that both low and high pixel values can be accurately recorded on the same frame: while the low-resolution spots may be strong enough to approach the upper limit of what a camera can measure, the high-resolution spots may be barely discernible over the background. Furthermore, under continuous rotation of the sample, the dead time during detector readout must be minimal, otherwise systematic gaps will be introduced in the sampling of reciprocal space.

While the majority of cryo-EMs these days are equipped with sensitive direct electron detectors for imaging, these cameras have not been used for MicroED because of concerns over damage to the sensor by the intense, incident beam. Instead, MicroED data have been collected on cameras that are not typically used for routine structure determination in other cryo-EM modalities, such as single particle analysis and cryotomograpy. As a result, the number of MicroED practitioners has been limited because most facilities do not have the resources to provide a dedicated camera for MicroED. If MicroED data can be collected using the very same direct electron detectors used for single particle analysis, the number of laboratories with the ability to conduct MicroED measurements could increase substantially.

We collected MicroED data from microcrystals of proteinase K using the Thermo Fisher Falcon III direct electron detector in integrating mode and compared the resulting structure to results obtained using diffraction-optimized cameras such as the CMOS-based Thermo Fisher CetaD. Unlike the regular Ceta camera, the CetaD is fitted with a thicker scintillator to better capture weak intensities of high-resolution Bragg spots. We demonstrate that reliable structure solution is possible from a typical direct electron detecting camera, and that these cameras may even offer some advantages over those specifically designed for diffraction experiments. In order to facilitate this work, we developed the necessary software tools to convert the collected images from the Falcon III and CetaD into images that can be processed in standard data reduction suites such as *DIALS* (Winter *et al.*, 2018), *MOSFLM* (Leslie and Powell, 2007), and *XDS* (Kabsch, 2010). This software has been made freely available via our website (https://cryoem.ucla.edu/MicroED).

## Results

We collected data from six crystals of proteinase K: three measured in integrating mode on the Falcon III, and three on the CetaD (Table 1). Both cameras were configured for 2×2 binning, yielding 2048×2048 px^2^ frames; for proteinase K diffracting to ~2 Å resolution, this results in a 4-fold reduction of the data volume without causing detrimental spot overlap. On the CetaD camera in particular, the thicker scintillator is intended to trade spatial resolution for increased sensitivity. This modification is tailored towards diffraction measurements, where sensitivity is more important than spatial resolution. It also means that unbinned data offers little advantage on the CetaD (Tinti *et al.*, 2018).

**Table 1:**
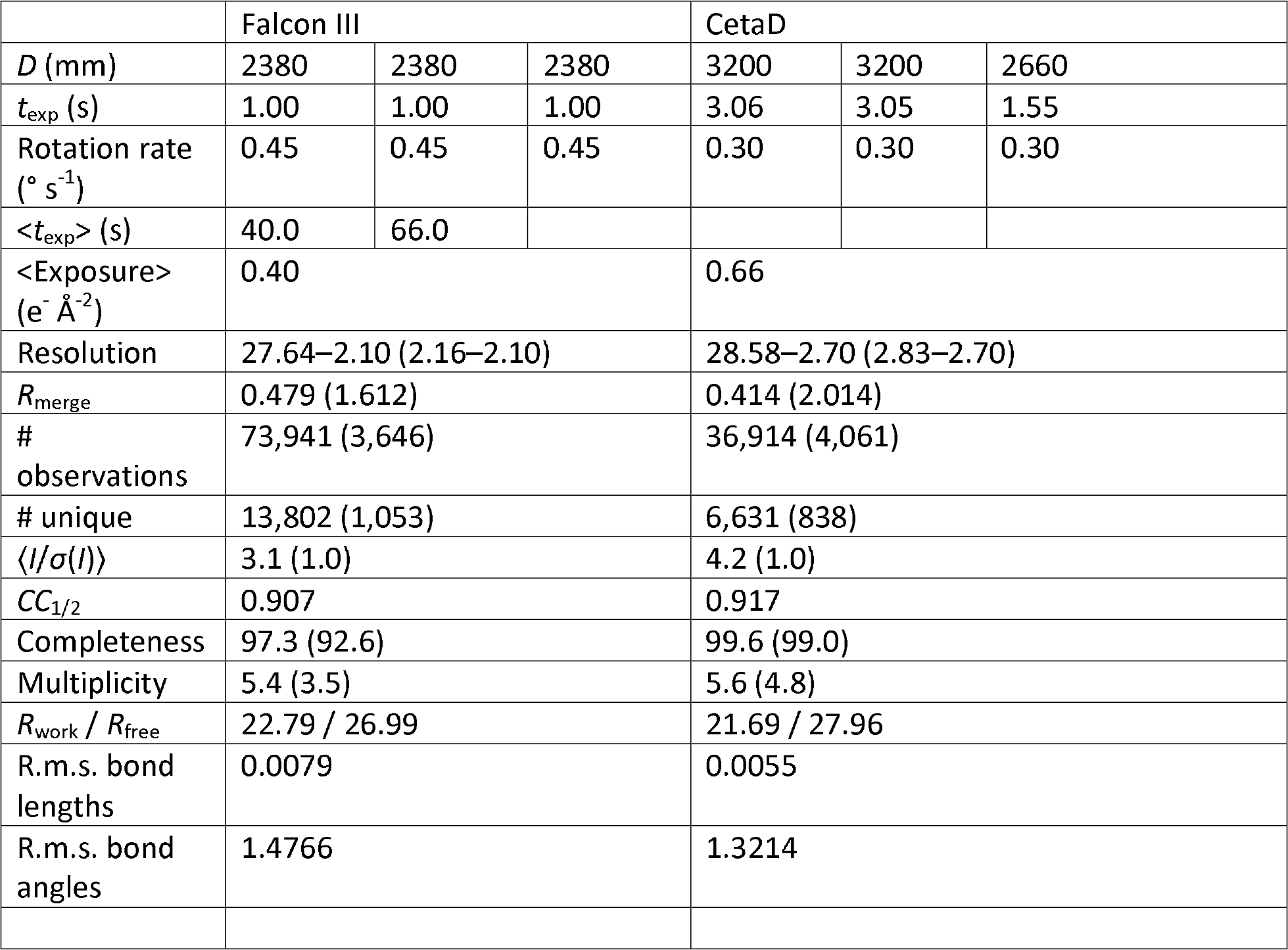
Processing and refinement statistics for proteinase K recorded on the Falcon III and CetaD cameras. D is the virtual sample–detector distance, which corresponds to the physical distance in an otherwise equivalent lensless system, and t_exp_ denotes the exposure time per frame during data collection. Note that not all collected frames were merged, and this is reflected in <t_exp_>, the mean cumulative irradiated time of all the frames in a multicrystal dataset. The average exposure, <Exposure>, is thus related to <t_exp_> by a multiplicative factor. Numbers in parentheses refer to the highest-resolution shell for merging. All data were collected at an acceleration voltage of 200 kV.

|  | Falcon III |  |  | CetaD |  |  |
| --- | --- | --- | --- | --- | --- | --- |
| $D$ (mm) | 2380 | 2380 | 2380 | 3200 | 3200 | 2660 |
| $t_{\text{exp}}$ (s) | 1.00 | 1.00 | 1.00 | 3.06 | 3.05 | 1.55 |
| Rotation rate<br>($^{\circ} \text{ s}^{-1}$) | 0.45 | 0.45 | 0.45 | 0.30 | 0.30 | 0.30 |
| $\langle t_{\text{exp}} \rangle$ (s) | 40.0 | 66.0 | | | | |
| $\langle \text{Exposure} \rangle$<br>($\text{e}^{-} \text{\AA}^{-2}$) | 0.40 | | | 0.66 | | |
| Resolution | 27.64–2.10 (2.16–2.10) |  |  | 28.58–2.70 (2.83–2.70) |  |  |
| $R_{\text{merge}}$ | 0.479 (1.612) | | | 0.414 (2.014) | | |
| # observations | 73,941 (3,646) |  |  | 36,914 (4,061) |  |  |
| # unique | 13,802 (1,053) |  |  | 6,631 (838) |  |  |
| $\langle I/\sigma(I) \rangle$ | 3.1 (1.0) | | | 4.2 (1.0) | | |
| $CC_{1/2}$ | 0.907 | | | 0.917 | | |
| Completeness | 97.3 (92.6) |  |  | 99.6 (99.0) |  |  |
| Multiplicity | 5.4 (3.5) |  |  | 5.6 (4.8) |  |  |
| $R_{\text{work}} / R_{\text{free}}$ | 22.79 / 26.99 | | | 21.69 / 27.96 | | |
| R.m.s. bond lengths | 0.0079 |  |  | 0.0055 |  |  |
| R.m.s. bond angles | 1.4766 |  |  | 1.3214 |  |  |

Using the Falcon III, 129 frames were collected from each crystal as 1 s exposures while continuously rotating the stage, whereas up to 71 frames were collected from CetaD. The exposure time for the crystals measured on the CetaD varied between 1.55 s and 3.06 s. In all cases the stage was rotated towards zero tilt, but the tilt rates were correspondingly higher on the Falcon III (0.45° s^−1^) than on the CetaD (0.30° s^−1^). Data were integrated to the edge of the detector (2.1 Å for Falcon III, 2.3–2.8 Å for CetaD), and all crystals were measured with an estimated exposure rate of 0.01 e^−^ Å^−2^ s^−1^ at an acceleration voltage of 200 kV. For comparison, data previously collected from proteinase K on a TVIPS TemCam-F416 under otherwise similar conditions (Hattne *et al.*, 2018) used significantly longer exposures (4–5 s), with correspondingly slower rotation rates (0.09° s^−1^)

With the Falcon III and the CetaD, data can be collected an order of magnitude faster compared to previous protocols for proteinase K (Hattne *et al.*, 2016, 2018; de la Cruz *et al.*, 2017). The increased sensitivity allows the per-frame exposure to be reduced during data collection because fewer electrons are required to obtain a sufficiently strong signal over the noise of the background. Combined with the faster readout rate, this implies that complete datasets can be collected both faster and using lower total exposure than previously possible. While the precise relationship between exposure and absorbed dose also depends on chemical composition, longer exposures always increase the absorbed dose. Since absorbed dose is directly related to radiation damage, the ability to obtain complete datasets with fewer e^−^ Å^−2^ is expected to translate to final models of proteins that are less damaged.

One of the first noticeable effects of radiation damage is the exponential falloff of intensity with increasing exposure (Blake and Phillips, 1962; Liebschner *et al.*, 2015; Hattne *et al.*, 2018), commonly characterized by the dose at which the average integrated intensity has dropped below some threshold. The rate of the intensity reduction is dependent on the sample, both its chemical composition and the size of the illuminated crystal (Nave and Hill, 2005), but should be unaffected by the detector. For proteinase K measured on the Falcon III and CetaD, the intensity has fallen to 50% of its extrapolated value at zero dose once exposed to 2.5 and 1.6 e^−^ Å^−2^, respectively (Figure 1(a)). These values agree with *D*_50_ = 2.2 e^−^ Å^−2^ previously found for the same sample measured on a TVIPS TemCam-F416 (Hattne *et al.*, 2018), and indicate that microscope and camera parameters are adequately calibrated.

**Figure 1:**
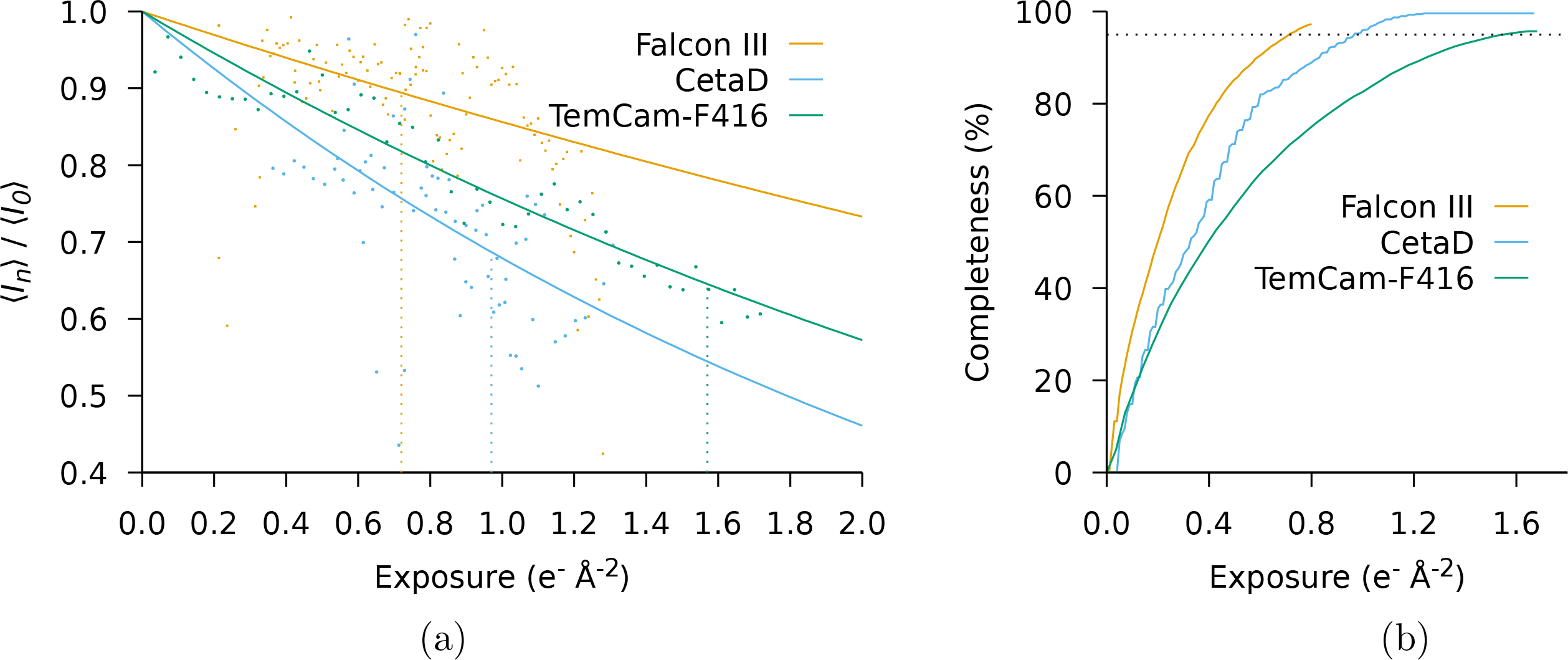
Mean intensity and completeness as a function of exposure. (a) For each camera, the integrated intensities on the subset of frames in the [−30°, +30°] tilt range were averaged; this reduces systematic effects on the intensities arising from longer paths through the sample at higher tilt. The reflections in the resolution range common to all datasets (20.0–2.70 Å) were then fit to a function on the form A_cryst_ exp(−B_cam_ × x), where A_cryst_ was refined for each crystal and B_cam_ was refined for each camera. The dotted vertical lines indicate the exposure at which 95% completeness was obtained. (b) The exposure-dependency of the completeness is determined by merging only frames with an average exposure less than the given value. The dotted horizontal line marks 95% completeness.

Figure 1(b) compares the completeness as a function of exposure. For three crystals, a total exposure of 0.97 e^−^ Å^−2^ frames was needed to reach 95% completeness on the CetaD, whereas only 0.72 e^−^ Å^−2^ were needed for the Falcon III. Both these exposures are significantly lower than the 1.57 e^−^ Å^−2^ previously required to reach the same completeness on the TVIPS TemCam-F416 (Hattne *et al.*, 2018).

The manifestation of damage in real space similarly agrees with previous observations. Figure 2 shows the density around the two disulfide bonds in proteinase K, where damage to the Cys 283-Cys 354 bond is immediately apparent in the data collected on the CetaD (total exposure 1.6 e^−^ Å^−2^). In comparison, the total exposure for the data collected on the Falcon III is about half (0.8 e^−^ Å^−2^). At this level of detail, damage to the disulfide bonds cannot be observed in the data collected on the Falcon III, even though the model was refined against higher resolution data, which is known to be affected by radiation damage earlier (Table 1).

**Figure 2:**
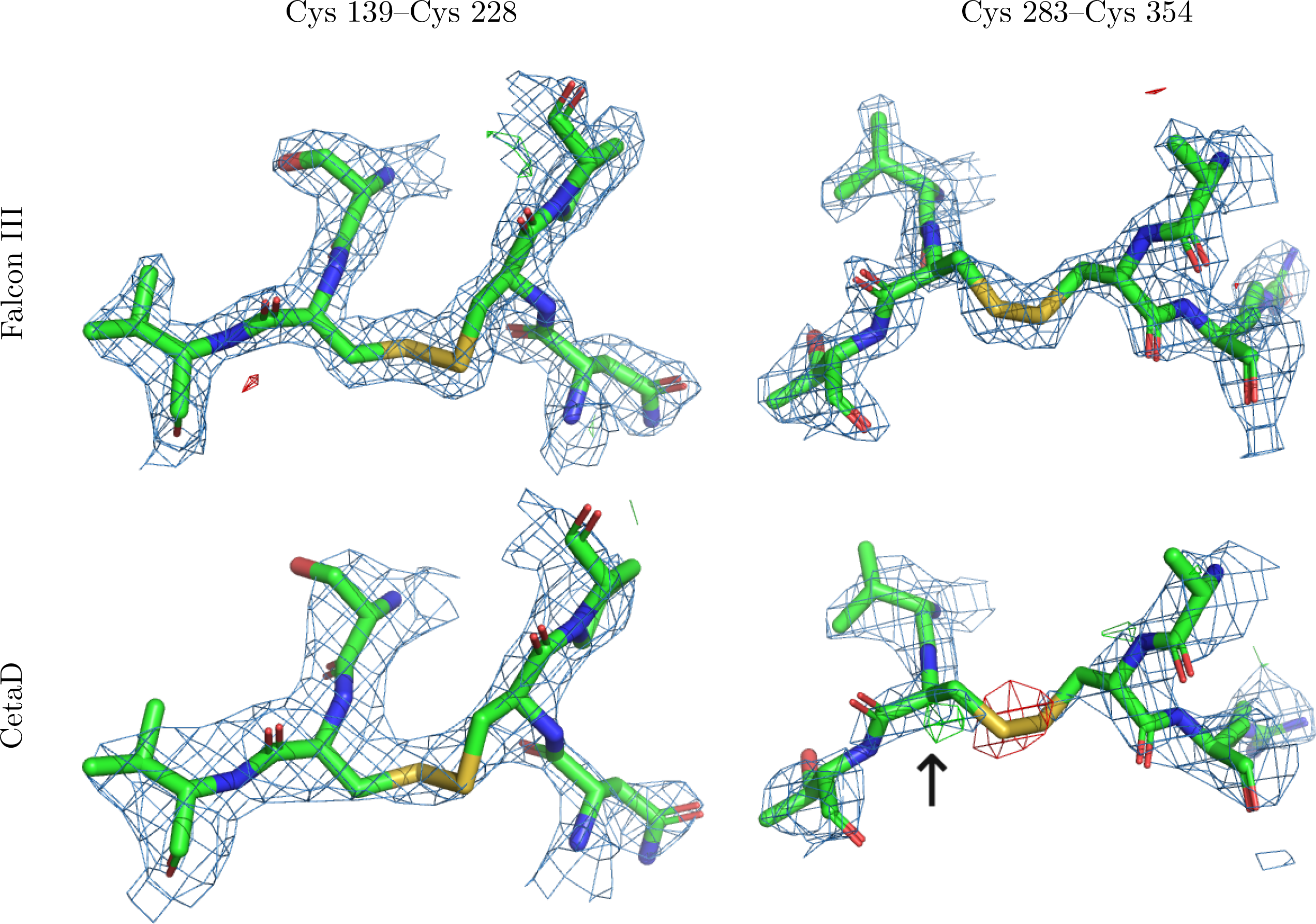
Density around the two disulfide bonds in proteinase K. The density around the two disulfide bonds indicates increasing radiation damage as an effect of increasing dose. The positive difference density around C_β_ of Cys 283 (black arrow) indicates a partially dislocated sulfur atom. The 2mF_o_-DF_c_ densities (blue meshes) are contoured at 1.5 σ above the mean, mFo-DFc difference densities (green/red meshes) are contoured at ±3 σ around the mean. All meshes were carved to 2 Å around the selected atoms in PyMol (Schrödinger LLC, 2014).

The modulation transfer function (MTF) at half Nyquist is higher on the Falcon III (0.25) than on the CetaD (0.06) (Kuijper *et al.*, 2015). Since the MTF is the modulus of the Fourier transform of the point-spread function (PSF), a higher MTF is expected to translate to sharper diffraction spots on the Falcon III. While the differences in spot size are barely discernible by visual inspection of the rocking curves (Figure 3), the differences are revealed by comparing the area of an asymmetric, two-dimensional Gaussian function fitted to the pixel values around each predicted spot location. The spot areas are weakly correlated with resolution, hence only spots in the resolution interval common to all datasets (20.0-2.7 Å) were considered. While the counts on the Falcon III are about an order of magnitude lower than on the CetaD, the average spot on the Falcon III is sharper than on the CetaD, and the area distribution is more peaked (Table 2, Figure 4).

**Table 2:**
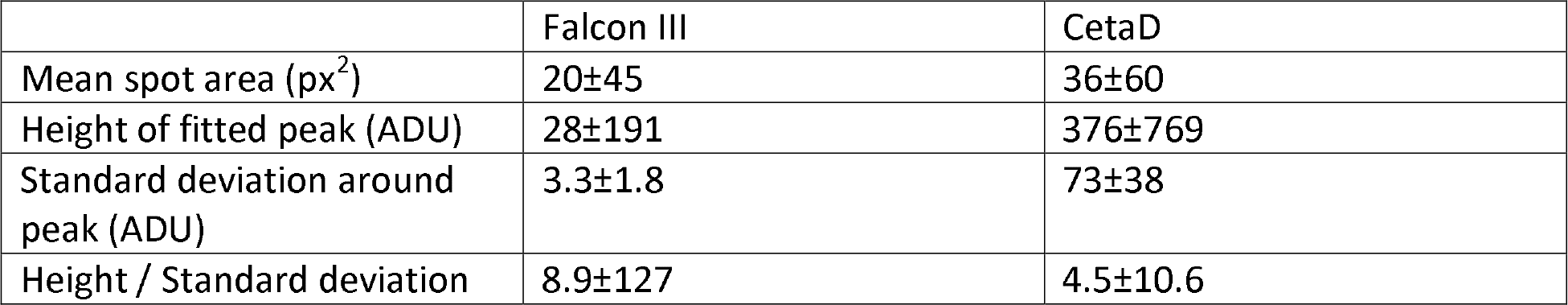
The ratio of peak height to the standard deviation in an annulus around the fitted peak is higher for the Falcon III than for the CetaD. The standard deviation was calculated from the pixel values in an elliptical annulus around the fitted peak, with inner diameters determined by the size of the fitted peak and outer diameters 4 px larger. Peaks were fitted as described in the caption of Figure 4.

|  | Falcon III | CetaD |
| --- | --- | --- |
| Mean spot area ($\text{px}^2$) | 20±45 | 36±60 |
| Height of fitted peak (ADU) | 28±191 | 376±769 |
| Standard deviation around peak (ADU) | 3.3±1.8 | 73±38 |
| Height / Standard deviation | 8.9±127 | 4.5±10.6 |

**Figure 3:**
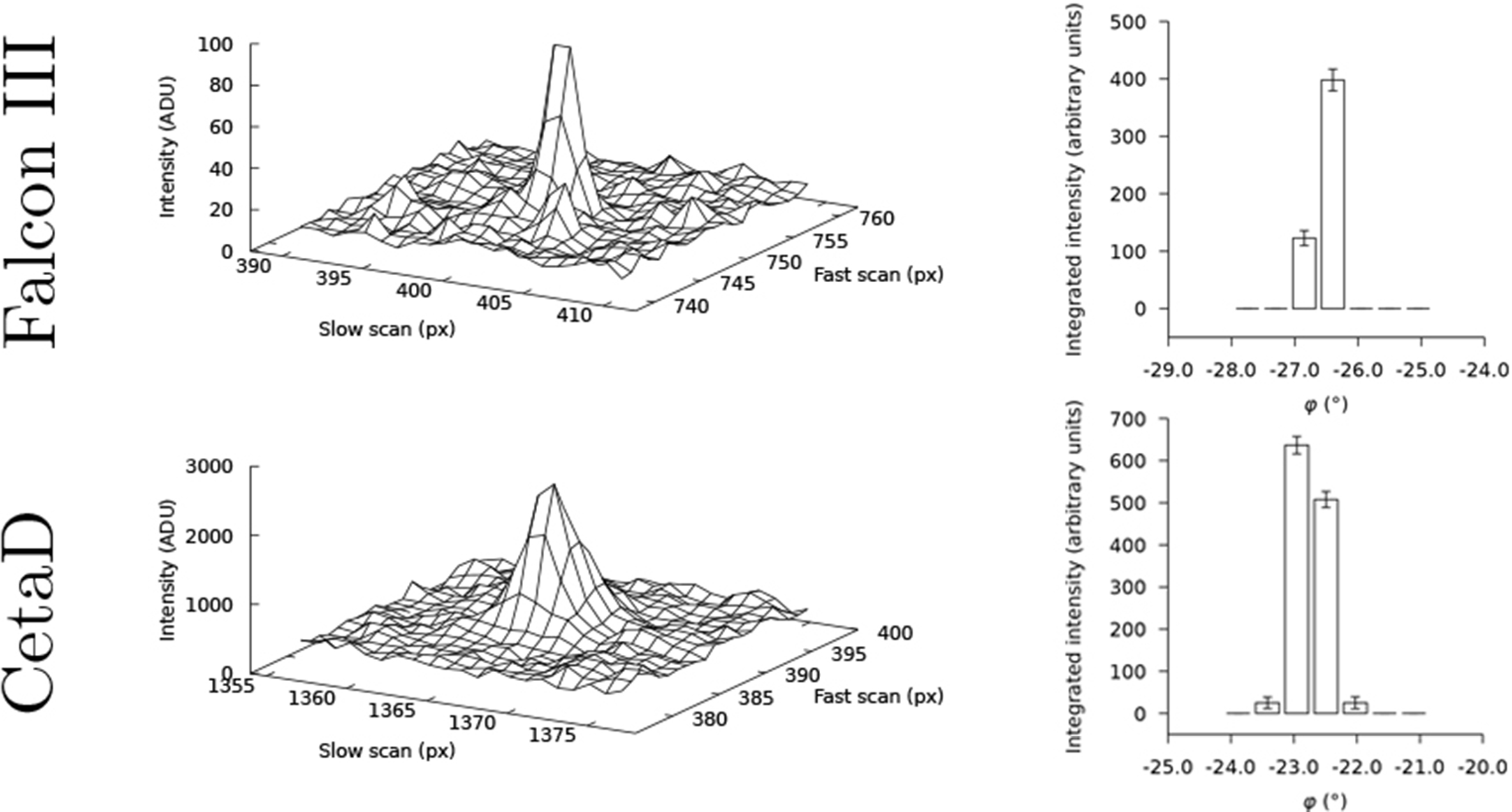
The (10, 16, 12) reflection at ~3.3 Å resolution on consecutive frames. Pixel values recorded on the Falcon III (top row; rotation range ∆φ = 0.45° per frame) and the CetaD (bottom row; rotation range ∆φ = 0.46° per frame) camera. The rightmost panel in each row shows the profile-fitted intensities as integrated by MOSFLM, where the error bars span one standard deviation. Constant pedestals were added in the conversion from the native camera format: the offset was 8 ADU for the Falcon III, and 128 ADU for the CetaD. The physical pixels sizes on both cameras are identical (14 μm, square). Note that the peak counts on the CetaD are more than an order of magnitude higher than on the Falcon III.

**Figure 4:**
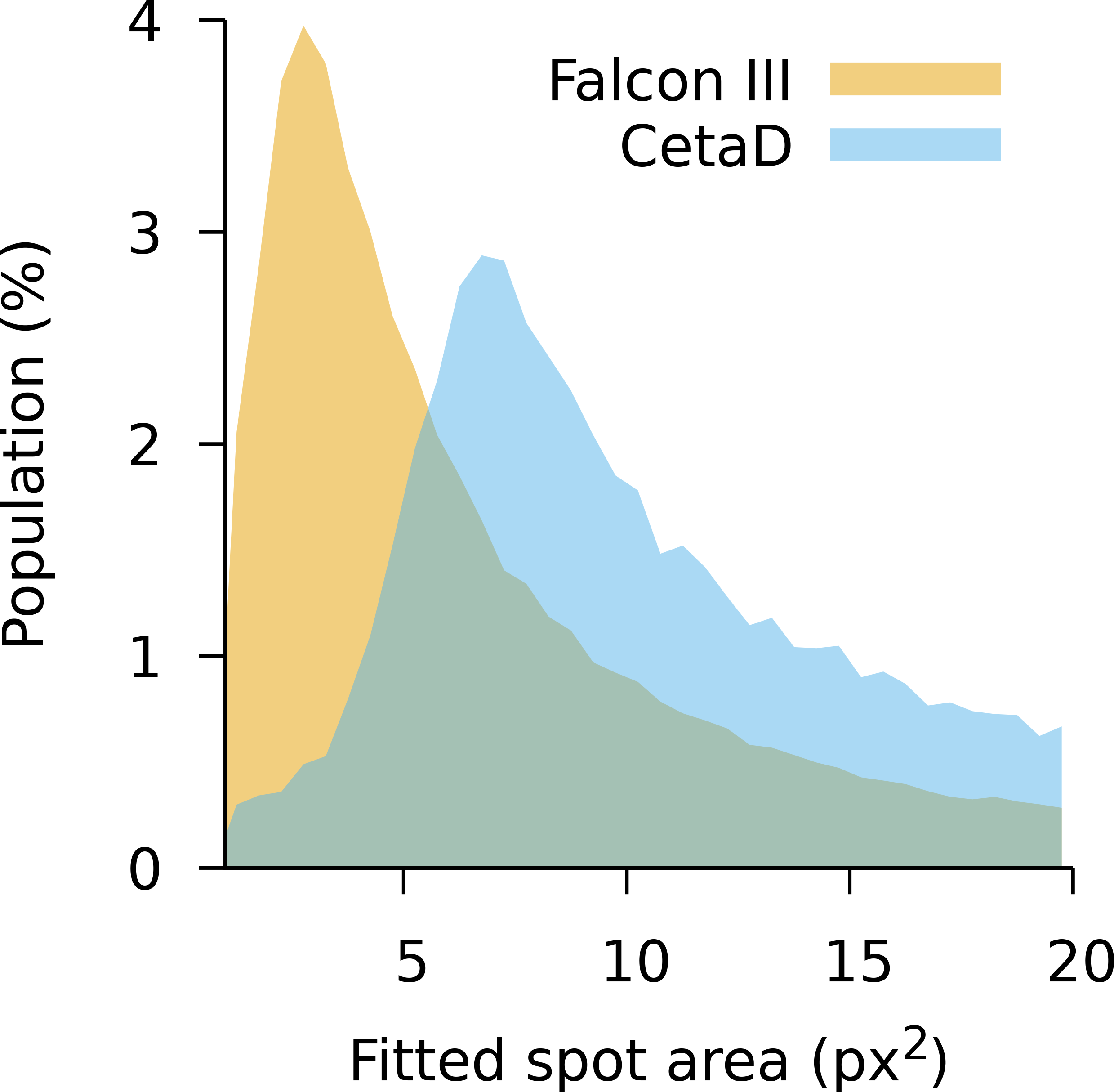
Histogram of the area of strong spots. The spot area, π σ_x_ σ_y_, determined by fitting a two-dimensional, elliptical Gaussian function A × exp(−**x**^T^ **R**^T^ **Σ**^−2^ **R x** / 2) + B, where **R** is a rotation matrix, **Σ** is a diagonal matrix of standard deviations σ_x_ and σ_y_, and A and B are scalars. All pixels in a 21×21 px^2^ box around the integrated spots for the Falcon III (orange) and CetaD (blue) cameras were considered in the fit. To control for the effect of resolution on the fitted spot sizes, only spots in the common resolution interval 20.0–2.7 Å resolution interval were considered. The counts for spots where the fit converged are scaled to equal area (N_Falcon III_ = 82,117 and N_CetaD_ = 68,654).

## Conclusion

The higher sensitivity and readout rate of the Falcon III and CetaD cameras allow complete datasets to be collected faster and with lower total exposure. The average time to record a single dataset from Proteinase K was less than half the time previously required to collect similar low-dose datasets (Hattne *et al.*, 2018). With the Falcon III and CetaD cameras, this can be accomplished without increasing the total dose deposited to the sample, thus keeping radiation damage minimal. In fact, the average exposure for the data merged from the Falcon III is also less than half of what was used for the TVIPS TemCam-F416.

Compared to the camera previously used to collect most of the MicroED data, both the Falcon III and the CetaD exhibit a higher modulation transfer function. This is true even for the CetaD with its thicker scintillator layer. A higher MTF, or equivalently, a narrower PSF, is expected to translate to a higher signal-to-noise ratio, as the diffraction peak is focused to a smaller area and becomes easier to distinguish over the background. It does, however, not necessarily imply a higher value of ⟨*I*/*σ*(*I*)⟩ (Table 1). Ideally, the mean ratio of the integrated intensity to its standard deviation is independent of the spot area, since all pixels in the peak are considered for the integrated intensity. In practice, the observed signal-to-noise ratio requires the per-pixel gain of the detector to be known to high accuracy (Leslie, 2006), which has been difficult to accomplish on the cameras used here. The signal-to-noise ratio is also affected by factors unrelated to the camera: for instance, the weak, high-resolution reflections will skew the distribution towards lower values for well-diffracting samples. Indeed, ⟨*I*/*σ*(*I*)⟩ is lower for the higher-resolution datasets collected on the Falcon III compared to the datasets measured on the CetaD (Table 1), even though the average peak height relative to the variance of the background is almost twice as high (Table 2).

The Falcon III camera implements an electron-counting mode in addition to the integrating mode used to collect the data here. Electron-counting has the potential for measuring data at near-optimal detective quantum efficiency (McMullan, Faruqi and Henderson, 2016), but requires that data collection is carefully designed to prevent saturating individual per-pixel counters. In particular, electron counting presents a formidable challenge for diffraction data, where the camera must accurately represent both high and low pixel counts. Since it is conceivable that the flux could be reduced, and the data collection correspondingly prolonged, future work is aimed to make electron-counting a feasible mode of data collection for MicroED.

Our results demonstrate that the typical direct electron detectors used for other cryo-EM modalities can also be used for MicroED alleviating the need for additional dedicated cameras. Compared to cameras used previously, the Falcon III and CetaD offer the possibility to collect complete data at lower exposure in a shorter amount of time. This has immediate implications for efforts to automate MicroED data collection (Jason de la Cruz *et al.*, 2019), where efficient use of shared resources may be a major concern, but also leads to structural models with limited or little radiation damage. For example, combining MicroED data collection in SerialEM (Jason de la Cruz *et al.*, 2019) with a Falcon III direct electron detector can result in more than 300 complete data sets collected autonomously overnight and this level of productivity is commensurate with X-ray crystallography at synchrotrons. As MicroED is gaining momentum in the cryoEM field that is already undergoing rapid changes, developing the MicroED data collection protocols and software analysis tools to optimally use new hardware will be a priority for the immediate future.

## Software availability

Software tools to convert the native output format, both MRC (Cheng *et al.*, 2015) and TIA series files, to SMV or TIFF, are available from https://cryoem.ucla.edu/MicroED and will be included in an upcoming release of the rebranded MicroED tools. The updated version also contains programs to directly convert data collected with SerialEM (Jason de la Cruz *et al.*, 2019).

## Methods

Proteinase K from *E. album* (Sigma-Aldrich, St Louis, MO, USA) was used without further purification to grow crystals in sittings drops (Hattne *et al.*, 2016). Protein powder dissolved in 50 mM Tris-HCl pH 8 was mixed with equal amounts of 1.25 M ammonium sulfate and dispensed in 24-well plates, where crystals with an average size of ~50 μm appeared in less than one hour. Sittings drops were diluted with well solution to a final volume of ~25 μl, and crystals, all from the same batch, were placed on glow-discharged Quantifoil R2/2 Cu300 grids by pipetting 2 μl onto the carbon side. After blotting from the back for 5 s at 4°C and 100% environment humidity, grids were vitrified and transferred to liquid nitrogen.

MicroED data were collected in an FEI Talos Arctica transmission electron microscope at an acceleration voltage of 200 kV. The temperature was maintained at <100 K while samples were continuously rotated in the electron beam and exposures, varying in duration between 1 and 3 s but constant for each crystal (Table 1), were collected on the different cameras.

Images were converted from the native output format of the camera to Super Marty View (SMV) format using software developed in house. The conversion programs extract the available metadata and automatically derive as much information as possible to allow downstream data reduction packages to reconstruct the diffraction geometry; parameters not contained in the output (*e.g*. the calibrated sample-detector distance) must be specified by the user during image conversion. Since negative pixel values are retained in the native format they need not be explicitly modelled (Hattne *et al.*, 2016), but are addressed by adding a user-determined, per-dataset constant to all pixel values of each frame (8 ADU for data from the Falcon III, 128 ADU for CetaD frames). Pixel values below this pedestal (≤0.03% per Falcon III frame, ≤0.003% for the CetaD) were set to zero and discarded during integration. No further corrections were applied; in particular, procedures to correct for drifting dark current were disabled.

Data were indexed and integrated in *MOSFLM* (Leslie and Powell, 2007) through its graphical interface *iMosflm* (Battye *et al.*, 2011), taking the previously added pedestal into account. The gain, *G*, was estimated assuming Poisson statistics of the background pixels, and held fixed for both the Falcon III (*G* = 1.0) and the CetaD (*G* = 14). After scaling and merging in *AIMLESS* (Evans and Murshudov, 2013), the data were phased by molecular replacement in *MOLREP* (Vagin and Teplyakov, 1997) using PDB ID 5k7s (de la Cruz *et al.*, 2017) as a search model. Atomic models were refined in *REFMAC* (Murshudov *et al.*, 2011); automated solvent modelling and manual curation was performed in *Coot* (Emsley *et al.*, 2010). Custom analysis tools were written in Python using the Computational Crystallography Toolbox (*cctbx*) (Grosse-Kunstleve *et al.*, 2002) and optimization routines implemented in SciPy (Oliphant, 2007).

## Acknowledgements

The Gonen laboratory is funded by the Howard Hughes Medical Institute. We thank Lingbo Yu (Thermo Fisher) for helpful discussions.

## References

Battye, T. G. G., Kontogiannis, L., Johnson, O., Powell, H. R. and Leslie, A. G. W. (2011) ‘iMOSFLM: a new graphical interface for diffraction-image processing with MOSFLM.’, Acta crystallographica. Section D, Biological crystallography. International Union of Crystallography, 67(Pt 4), pp. 271–81. doi:10.1107/S0907444910048675.

Blake, C. C. F. and Phillips, D. C. (1962) ‘Effects of X-irradiation on single crystals of myoglobin’. The Royal Institution, London, England., pp. 183–191.

Cheng, A., Henderson, R., Mastronarde, D., Ludtke, S. J., Schoenmakers, R. H. M., Short, J., Marabini, R., Dallakyan, S., Agard, D. and Winn, M. (2015) ‘MRC2014: Extensions to the MRC format header for electron cryo-microscopy and tomography’, Journal of Structural Biology. Elsevier Inc., 192(2), pp. 146–150. doi:10.1016/j.jsb.2015.04.002.

Clabbers, M. T. B., Gruene, T., Parkhurst, J. M., Abrahams, J. P. and Waterman, D. G. (2018) ‘Electron diffraction data processing with DIALS’, Acta Crystallographica Section D Structural Biology. International Union of Crystallography, 74(6), pp. 506–518. doi:10.1107/S2059798318007726.

Emsley, P., Lohkamp, B., Scott, W. G. and Cowtan, K. (2010) ‘Features and development of Coot.’, Acta crystallographica. Section D, Biological crystallography. International Union of Crystallography, 66(Pt 4), pp. 486–501. doi:10.1107/S0907444910007493.

Evans, P. R. and Murshudov, G. N. (2013) ‘How good are my data and what is the resolution?’, Acta Crystallographica Section D Biological Crystallography. International Union of Crystallography, 69(7), pp. 1204–1214. doi:10.1107/S0907444913000061.

Grosse-Kunstleve, R. W., Sauter, N. K., Moriarty, N. W. and Adams, P. D. (2002) ‘The Computational Crystallography Toolbox: crystallographic algorithms in a reusable software framework’, Journal of Applied Crystallography. International Union of Crystallography, 35(1), pp. 126–136. doi:10.1107/S0021889801017824.

Hattne, J., Shi, D., Glynn, C., Zee, C., Gallagher-Jones, M., Martynowycz, M. W., Rodriguez, J. A. and Gonen, T. (2018) ‘Analysis of Global and Site-Specific Radiation Damage in Cryo-EM’, Structure. Elsevier, 26(5), p. 759–766.e4. doi:10.1016/j.str.2018.03.021.

Hattne, J., Shi, D., de la Cruz, M. J., Reyes, F. E. and Gonen, T. (2016) ‘Modeling truncated pixel values of faint reflections in MicroED images’, Journal of Applied Crystallography, 49(3), pp. 1029–1034. doi:10.1107/S1600576716007196.

Henderson, R. (1995) ‘The potential and limitations of neutrons, electrons and X-rays for atomic resolution microscopy of unstained biological molecules’, Quarterly Reviews of Biophysics, 28(02), p. 171. doi:10.1017/S003358350000305X.

Jason de la Cruz, M., Martynowycz, M. W., Hattne, J. and Gonen, T. (2019) ‘MicroED data collection with SerialEM’, Ultramicroscopy. Elsevier B.V., 201(January), pp. 1–4. doi:10.1016/j.ultramic.2019.03.009.

Jones, C. G., Martynowycz, M. W., Hattne, J., Fulton, T. J., Stoltz, B. M., Rodriguez, J. A., Nelson, H. M. and Gonen, T. (2018) ‘The CryoEM Method MicroED as a Powerful Tool for Small Molecule Structure Determination’, ACS Central Science. American Chemical Society, 4(11), pp. 1587–1592. doi:10.1021/acscentsci.8b00760.

Kabsch, W. (2010) ‘XDS.’, Acta crystallographica. Section D, Biological crystallography, 66(Pt 2), pp. 125–32. doi:10.1107/S0907444909047337.

Kuijper, M., van Hoften, G., Janssen, B., Geurink, R., De Carlo, S., Vos, M., van Duinen, G., van Haeringen, B. and Storms, M. (2015) ‘FEI’s direct electron detector developments: Embarking on a revolution in cryo-TEM’, Journal of Structural Biology. Elsevier Inc., 192(2), pp. 179–187. doi:10.1016/j.jsb.2015.09.014.

de la Cruz, M. J., Hattne, J., Shi, D., Seidler, P., Rodriguez, J., Reyes, F. E., Sawaya, M. R., Cascio, D., Weiss, S. C., Kim, S. K., Hinck, C. S., Hinck, A. P., Calero, G., Eisenberg, D. and Gonen, T. (2017) ‘Atomic-resolution structures from fragmented protein crystals with the cryoEM method MicroED’, Nature Methods. Nature Publishing Group, 14(4), pp. 399–402. doi:10.1038/nmeth.4178.

Leslie, A. G. W. (2006) International Tables for Crystallography, International Tables for Crystallography Volume F: Crystallography of biological macromolecules. Edited by M. G. Rossmann and E. Arnold. Chester, England: International Union of Crystallography (International Tables for Crystallography). doi:10.1107/97809553602060000106.

Leslie, A. G. W. and Powell, H. R. (2007) ‘Processing diffraction data with mosflm’, in Read, R. J. and Sussman, J. L. (eds) Evolving Methods for Macromolecular Crystallography. Dordrecht: Springer Netherlands (NATO Science Series II: Mathematics, Physics and Chemistry), pp. 41–51. doi:10.1007/978-1-4020-6316-9_4.

Liebschner, D., Rosenbaum, G., Dauter, M. and Dauter, Z. (2015) ‘Radiation decay of thaumatin crystals at three X-ray energies’, Acta Crystallographica Section D: Biological Crystallography. International Union of Crystallography, 71, pp. 772–778. doi:10.1107/S1399004715001030.

McMullan, G., Faruqi, A. R. and Henderson, R. (2016) ‘Direct Electron Detectors’, in The Resolution Revolution: Recent Advances In cryoEM. 1st edn. Elsevier Inc., pp. 1–17. doi:10.1016/bs.mie.2016.05.056.

Murshudov, G. N., Skubák, P., Lebedev, A. a, Pannu, N. S., Steiner, R. a, Nicholls, R. a, Winn, M. D., Long, F. and Vagin, A. a (2011) ‘REFMAC5 for the refinement of macromolecular crystal structures.’, Acta crystallographica. Section D, Biological crystallography, 67(Pt 4), pp. 355–67. doi:10.1107/S0907444911001314.

Nannenga, B. L., Shi, D., Leslie, A. G. W. and Gonen, T. (2014) ‘High-resolution structure determination by continuous-rotation data collection in MicroED’, Nature Methods, 11(9), pp. 927–930. doi:10.1038/nmeth.3043.

Nave, C. and Hill, M. A. (2005) ‘Will reduced radiation damage occur with very small crystals?’, Journal of synchrotron radiation, 12(Pt 3), pp. 299–303. doi:10.1107/S0909049505003274.

Oliphant, T. E. (2007) ‘Python for Scientific Computing’, Computing in Science & Engineering, 9(3), pp. 10–20. doi:10.1109/MCSE.2007.58.

Pflugrath, J. W. (1999) ‘The finer things in X-ray diffraction data collection’, Acta Crystallographica Section D Biological Crystallography, 55(10), pp. 1718–1725. doi:10.1107/S090744499900935X.

Rodriguez, J. A., Ivanova, M. I., Sawaya, M. R., Cascio, D., Reyes, F. E., Shi, D., Sangwan, S., Guenther, E. L., Johnson, L. M., Zhang, M., Jiang, L., Arbing, M. A., Nannenga, B. L., Hattne, J., Whitelegge, J., Brewster, A. S., Messerschmidt, M., Boutet, S., Sauter, N. K., Gonen, T. and Eisenberg, D. S. (2015) ‘Structure of the toxic core of α-synuclein from invisible crystals’, Nature, 525(7570), pp. 486–490. doi:10.1038/nature15368.

Schrödinger LLC (2014) ‘The PyMOL Molecular Graphics System’. Available at: https://www.pymol.org.

Shi, D., Nannenga, B. L., Iadanza, M. G. and Gonen, T. (2013) ‘Three-dimensional electron crystallography of protein microcrystals’, eLife, 2, p. e01345. doi:10.7554/eLife.01345.

Tinti, G., Fröjdh, E., van Genderen, E., Gruene, T., Schmitt, B., de Winter, D. A. M., Weckhuysen, B. M. and Abrahams, J. P. (2018) ‘Electron crystallography with the EIGER detector’, IUCrJ, 5(2), pp. 190–199. doi:10.1107/S2052252518000945.

Vagin, A. and Teplyakov, A. (1997) ‘MOLREP⍰: an Automated Program for Molecular Replacement’, Journal of Applied Crystallography. International Union of Crystallography, 30(6), pp. 1022–1025. doi:10.1107/S0021889897006766.

Winter, G., Waterman, D. G., Parkhurst, J. M., Brewster, A. S., Gildea, R. J., Gerstel, M., Fuentes-Montero, L., Vollmar, M., Michels-Clark, T., Young, I. D., Sauter, N. K. and Evans, G. (2018) ‘DIALS⍰: implementation and evaluation of a new integration package’, Acta Crystallographica Section D Structural Biology. International Union of Crystallography, 74(2), pp. 85–97. doi:10.1107/S2059798317017235.

